# Postprocessing of P300 Speller Output with a Large Language Model

**DOI:** 10.64898/2026.06.24.734268

**Authors:** Nikita Paplavsky, Mikhail Lebedev

## Abstract

P300 spellers convert electroencephalographic (EEG) activity into text by presenting users with a matrix of flickering characters. While these systems can achieve high classification accuracy, communication is severely slowed by the need for many stimulus repetitions to obtain a reliable signal. Reducing repetitions accelerates spelling but introduces character-level errors: insertions, deletions, and substitutions that degrade usability and increase user fatigue. Although substantial research has focused on improving performance at the signal acquisition and decoding stages, here we investigate a complementary text post-processing approach that leverages large language models (LLMs) to restore corrupted P300 speller output. We constructed a dataset derived from cLang-8 and simulated realistic P300-style text corruption using both random and empirically derived human-like error strategies. We evaluated several instruction-tuned LLMs alongside an optical character recognition (OCR)-fine-tuned ByT5 model under zero-shot and few-shot prompting conditions. We found that LLMs effectively recovered clean text from noisy inputs. Models employing SentencePiece tokenization consistently outperformed byte-pair encoding (BPE)-based counterparts, and few-shot in-context learning further improved restoration accuracy, with Gemma 3 achieving the strongest performance across all settings. These results suggest that LLM-based post-processing could enable P300 speller systems to operate with fewer repetitions and lower latency while maintaining or improving output accuracy, offering a practical path toward more efficient daily communication for users with motor and speech disabilities.

## Introduction

Brain-computer interfaces (BCI) that translate neural signals into text, commonly known as BCI spellers, offer a communication pathway for individuals with severe motor and speech impairments. Foundational studies using both intracranial (Kennedy and Bakay, 1998) and EEG-based (Birbaumer et al., 1999) systems established that even completely paralyzed patients can control computer applications through neural modulation. Among the various speller paradigms, the P300 speller introduced by Farwell and Donchin (1988) remains a benchmark due to its ease of implementation and dependable accuracy, though its information transfer rate is limited to 1–2 words per minute. In its conventional form, this system presents a character matrix in which rows and columns flash in random order; the user attends to their desired character, and the system decodes this selection by identifying the ensuing P300 event-related potential. Subsequent research has substantially enhanced the original design, boosting both speed and precision through improved signal processing, adaptive visual interfaces, and advanced classifiers (Fazel-Rezai et al., 2012; Krusienski et al., 2006; Rezeika et al., 2018). These refinements have translated into meaningful clinical applications, with proven efficacy for patients with amyotrophic lateral sclerosis (Sellers and Donchin, 2006; Nijboer et al., 2008; Guy et al., 2018) and those with aphasia resulting from stroke (Kleih and Botrel, 2024).

Despite years of development, P300 spellers still face substantial challenges. The generated text often contains errors stemming from the low signal-to-noise ratio of EEG recordings and the inherent variability in users’ cognitive and attentional states. These errors manifest as insertions, deletions, or substitutions in the decoded character sequence, undermining system usability. Several strategies have been proposed to mitigate these errors. One approach focuses on modifying the visual stimulation paradigm to boost the amplitude and consistency of the P300 response (Fazel-Rezai 2007; Allison and Pineda 2003; Guger et al. 2009), for instance by altering how symbols are flashed, their spatial layout, timing, and visual properties. A second line of work targets EEG signal processing, with signal averaging over trials being the most prevalent technique: multiple repetitions of the stimuli are collected and averaged to suppress noise. While this improves classification accuracy, it does so at the cost of additional time, which reduces the information transfer rate. Every extra repetition makes everyday communication more fatiguing. This reveals a fundamental trade-off: improving reliability through more repetitions directly conflicts with the practical need for fast, efficient communication.

To partially relax this trade-off, an attractive alternative is to apply post-processing not to the raw EEG signals, but to the already decoded text (Caria 2025; Mora-Cortes et al. 2014; Speier et al. 2016). If such methods can reliably correct typical spelling errors, the need for multiple signal repetitions could be reduced, potentially increasing throughput without sacrificing accuracy. In our recent work, we introduced a framework that harnesses the advanced contextual reasoning of large language models (LLMs) to address the inaccuracies inherent in BCI signal interpretation (Lebedev et al. 2025). Applying this framework to a P300 speller dataset, we modeled a communication process in which users construct words letter by letter, producing text with characteristic recognition errors. An LLM then refines this output by identifying and correcting these mistakes. Our results demonstrate that this integration can boost communication speed by reducing reliance on perfect single-character classification. This study added to the growing body of work on using language models to augment BCI communication (Caria 2025; Mora-Cortes et al. 2014; Speier et al. 2016), which spans from early systems employing dictionary-driven word prediction (Kaufmann et al. 2012; Lee et al. 2011; Ryan et al. 2010) to more recent platforms, such as ChatBCI (Hong et al. 2024) and Mind-Chat (Wang et al. 2025), that integrate advanced LLMs into P300 spellers to propose and anticipate words.

The idea of refining imperfectly reconstructed text through post-processing has deep roots in other domains, most notably optical character recognition (OCR). In OCR, visual ambiguity or degraded image quality often leads to uncertain character-level predictions (Do et al., 2025), creating a critical need for language-driven post-processing to polish raw outputs, correct improbable sequences, and restore coherent text. Building on this foundation, the recent rise of large language models (LLMs) has introduced substantially more powerful correction frameworks for OCR-like tasks. These frameworks accept noisy or partially corrupted input and harness the model’s linguistic knowledge to recover the most semantically and syntactically plausible original sentence.

While it is evident that integrating an LLM into a P300 speller has the potential to significantly enhance overall performance, a rigorous quantitative evaluation is essential to truly understand the bounds of this improvement. Specifically, it is critical to determine the maximum allowable error rate in the speller-generated output, that is, how many character-level or word-level mistakes the system can produce while still enabling the LLM to reliably infer and reconstruct the user’s intended message. Establishing this error tolerance threshold will not only clarify the conditions under which the LLM adds value, but also guide the development of more robust correction mechanisms. Furthermore, not all LLMs are created equal; a comparative analysis of multiple models is necessary to identify the optimal trade-off between correction accuracy, inference speed, and model size, especially for real-time or resource-constrained BCI applications. By systematically benchmarking different LLMs, we can pin-point the most suitable candidate that balances performance with deployability. With such quantitative insights, developers will be better positioned to co-design the P300 speller and the language model in tandem, tailoring signal processing, classification thresholds, and interaction paradigms to complement the strengths of the chosen LLM.

In this study, we evaluated the ability of a LLM to reconstruct original text from noisy character sequences that emulate realistic decoding errors produced by a P300 speller. We specifically focused on errors of human origin and adjacency errors (Fazel-Rezai R., 2007), where a character is mistakenly replaced by a neighboring character in the spelling grid or layout, as these error types are particularly characteristic of P300-based communication paradigms. We investigated how restoration quality depends on the length of the original text, given that P300 spelling sessions can generate messages of widely varying sizes, from isolated words to longer phrases and full sentences. Accordingly, we examined the capabilities of an LLM-based post-processing strategy to recover clean text from inputs corrupted by human and adjacency errors, and we systematically analyzed the relationship between input length and correction performance. To this end, we constructed noisy inputs by introducing controlled error patterns into target sentences and then prompted an LLM to predict the corrected versions. The outputs were evaluated against the ground-truth text using standard text similarity metrics and error statistics. The overall pipeline is illustrated in Fig. 1. We found that LLM-based correction remained highly effective under both random and human-like corruption, with the Gemma 3 model achieving the strongest overall performance. Notably, SentencePiece-based models consistently outperformed their BPE-based counterparts, and human-like corruption proved slightly more challenging than random errors; nevertheless, both conditions yielded robust restoration performance across the tested message lengths.

**Fig. 1.**
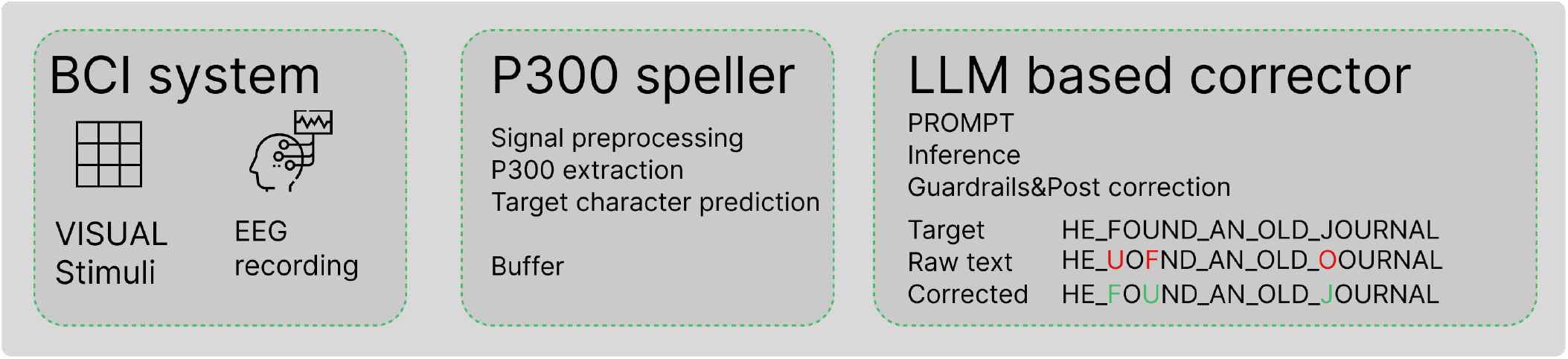
LLM-driven pipeline for EEG signal correction. The system acquires neural signals, translates them into text via a P300 speller, and utilizes an LLM to correct decoding errors, producing an accurate final transcription.

Overall, by framing P300-speller post-processing as text correction, we leverage LLMs as principled, context-sensitive inference engines to complement traditional EEG classification. Our results show that this synergy is transformative: LLM-based post-processing can recover coherent text from degraded EEG predictions, enabling robust performance with up to 50% fewer stimulus repetitions and lower latency without sacrificing accuracy relative to high-repetition baselines. These findings establish LLM-enhanced post-processing as a practical, deployable paradigm for BCIs, repositioning language models from an auxiliary tool to a core architectural pillar in P300-speller design for more responsive, human-centric communication.

## Materials and Methods

### Problem formulation

The goal of this study was to estimate the quality of text restoration using an LLM where the input text is corrupted by character-level errors that mimic the P300 speller output. We formulated this as a text restoration accuracy problem. Let *S* = {*s*_1_, … , *s*_*N*_} be a set of sentences, where each sentence *s*_*i*_ is a sequence of characters. For each character in each sentence, we apply a corruption process with a global error probability *p* ∈ [0, 1]. With probability *p*, a character is replaced according to a given error model; with probability 1 − *p*, it is left unchanged. The LLM then receives the corrupted sentence and produces a corrected version. Our aim is to characterize the relationship between the probability of error *p* and the ability of the LLM to restore the original text.

### Dataset and augmentations

The original 6 × 6 character matrix proposed by Farwell and Donchin, shown in Fig. 2, contains 36 symbols: the letters A–Z, the digits 0–9, and the underscore. This restricted symbol inventory imposes specific constraints on dataset design and makes it necessary to use a task-specific corpus for evaluation. In addition, the dataset should reflect non-professional lexicon to better match the intended correction setting. Since no suitable open dataset was available, we prepared one for this study.

**Fig. 2.**
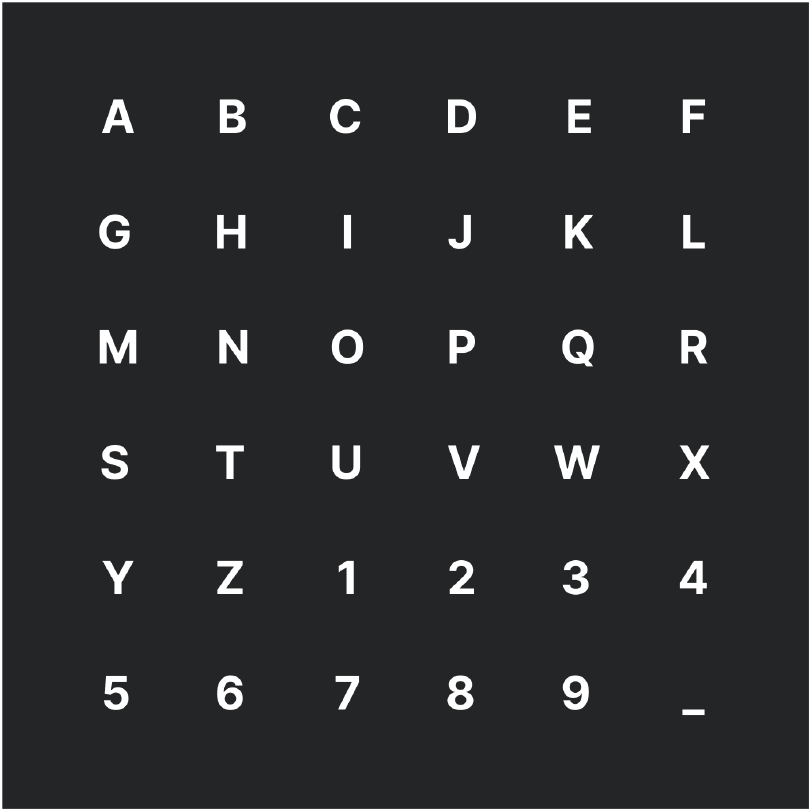
Original 6 × 6 character grid proposed by Farwell and Donchin

As a base dataset we chose cLang-8 dataset, following (Rothe et al. 2021), because it is a cleaned version of the widely used Lang-8 learner corpus and therefore provides non-professional writings well suited for correction experiments. Lang-8 itself was originally collected from a language-learning social network, where learner sentences were paired with human corrections, making it a natural source of clean text for grammatical and text-correction tasks (Mizumoto et al., 2011; Rothe et al., 2021). For our setting, this choice was especially appropriate because the target task was post-correction of decoding errors rather than correction of formally edited text. We prepared a task-specific version of the dataset by filtering out lines containing numbers and non-ASCII characters. After filtering, we normalized the text by expanding contractions, removing symbols that were not present in the original 6×6 character matrix (Fig. 2) and converting all text to lowercase. This processing step was designed to align the data with the symbol inventory available in the brain–computer interface setup. As a result, the final corpus contained clean, normalized examples that were consistent with the experimental decoding space. This pre-processing madee more homogeneous by reducing the variations unrelated to the correction task. The resulting dataset served as a controlled benchmark for evaluating LLM-based post-correction methods.

To simulate decoding errors typical of P300 spellers, we employed two complementary text corruption strategies. In the first strategy, we assumed that characters were selected from a 6 × 6 spelling matrix (Fig. 2), as in the standard row– column P300 speller layout. For each character in a sentence, with probability *p* we replaced the original character by sampling a random symbol from the same 6 × 6 matrix, and with probability 1 − *p* we kept the character unchanged. This produced corrupted sentences that reflected random misselections within the fixed symbol set used by the speller. The exact composition of the 6 × 6 matrix followed the original matrix design used in the P300 paradigm

The second strategy was designed to mimic realistic human error patterns in row–column P300 paradigms. This strategy was based on empirical data reported in (Gavett S. et. al 2012), which described typical confusion patterns between target and non-target symbols in a row–column P300 speller. We constructed an augmentation procedure that reproduced human errors in the RC paradigm using these data. Based on empirical data on error distribution, we created a normalized, weighted sampling distribution. Then, for each character in the set, we created an individual error set by shifting the distribution to the current character’s position. Specifically, we used the reported error statistics to define a distribution of likely corrupted outputs for each target symbol. We then sampled substitutions from this distribution when applying errors with probability *p*.

For both strategies, we generated multiple corrupted versions of each sentence across a range of error probabilities. Each corrupted set contained 500 sentences with lengths varying from 5 to 15 words. We used 12 corruption levels per strategy, with error probability *p* ranging from 0.025 to 0.5.

### Text correction approaches

Next, we explore three LLM approaches: a vanilla LLM-based method with two inference settings (zero-shot and 3-shot prompting) and an OCR correction model based on ByT5. For the n-shot LLM-based approach, we selected three compact yet effective instruction-tuned models: Gemma 3 4B IT, Qwen3 4B Instruct, and Llama 3.2 4B Instruct. We chose these models because they offer a favorable balance between performance, open availability, and near real-time inference on a single GPU, with inference time of less than one second per sentence.

### Zero shot inference

For zero-shot inference, we used a single prompt without providing any additional text restoration examples. This setting allowed the model to perform correction solely on the basis of the input sentence and the task instruction, without in-context demonstrations.

### 3-shot inference

For 3-shot inference, we prepended three exemplars of corrupted–corrected text pairs to the target input. This minimal in-context learning setup provided the model with sufficient demonstrations to perform the restoration task while maintaining a concise prompt.

### OCR fine-tuned ByT5 model for error correction

In the third approach, we employed a ByT5-based model fine-tuned specifically for OCR correction. ByT5 is a tokenizer-free variant of Google’s T5 architecture and closely follows the design of mT5. Pre-trained solely on mC4 using span masking over UTF-8 character sequences—and without any supervised objective—the model requires fine-tuning to be effectively applied to downstream tasks. This made it particularly well suited for noisy text restoration, where character-level distortions are frequent. In our setup, the model received only the corrupted text as input, with no additional prompts beyond the instruction to restore the sentence.

### Implementation

All experiments were conducted on a single consumer-grade NVIDIA RTX 4080 GPU with 16 GB of VRAM. The implementation was developed in Python using the Hugging Face Transformers library. Each test set contained 500 sentences. Depending on the model, inference time ranged from 0.8 to 2.2 seconds per sentence.

### Evaluation metrics

We used two complementary metrics to quantify the quality of text restoration: average word restoration accuracy and semantic textual similarity.

Average word restoration accuracy was defined as the ratio of correctly restored words to the total number of corrupted words in set. Formally,

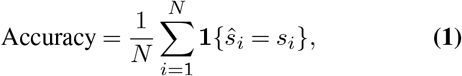

where *s*_*i*_ is the original word, *ŝ*_*i*_ is the LLM restored word, and **1**{·} is the indicator function.

Semantic textual similarity (STS) was measured by comparing the embeddings of the original and restored sentences. For each pair (*s*_*i*_, *ŝ*_*i*_), we computed sentence embeddings and then derived a similarity score from the distance between them in the embedding space. Higher scores indicated that the restored sentence was semantically closer to the original, even when character-level differences were present. For this purpose, we used the `sup-simcse-roberta-large` model (Tianyu Gao et. al 2021). This model was fine-tuned on natural language inference (NLI) data, which allowed it to better capture logical contradictions compared with conventional embedding models such as BERT.

## Results

For each corruption strategy and each error probability *p*, we computed the average word restoration accuracy and the average STS score over all sentences in the evaluation set. Figures 3 and 4 illustrate the dependency of accuracy and STS score on the error probability for the random 6 × 6 corruption. Figures 5 and 6 show these dependencies for the human-like error scenario based on row-column paradigm augmentation, which mimics realistic error distributions. These results enable comparison of the robustness of LLM-based correction across different error regimes and allow assessment of the sensitivity of restoration performance to the level and type of character-level corruption in the corrupted text.

**Fig. 3.**
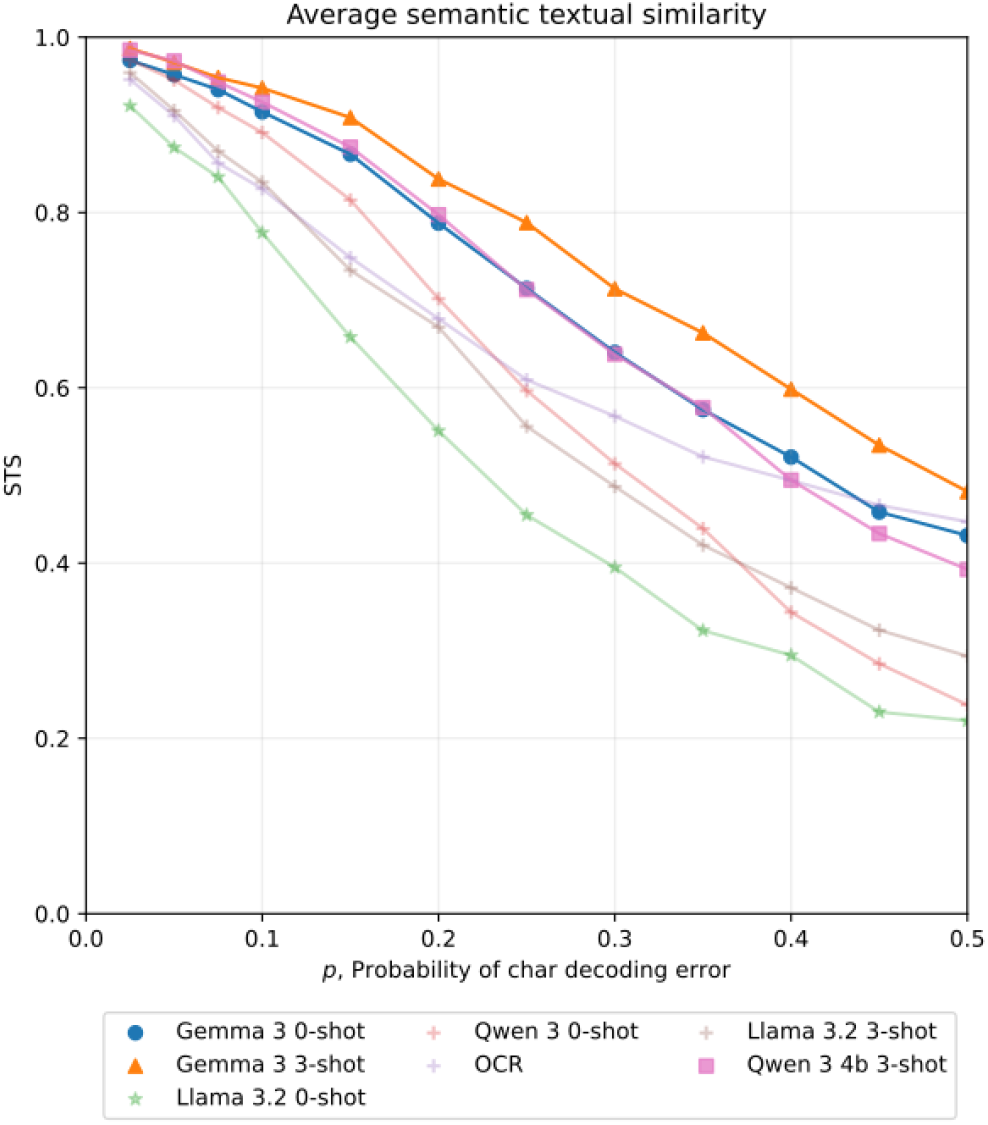
Average sematic textual similarity score (STS) for the reconstructed text as the function of spelling error probability for random errors. Curves for several LLMs are shown. (See key at the bottom.)

**Fig. 4.**
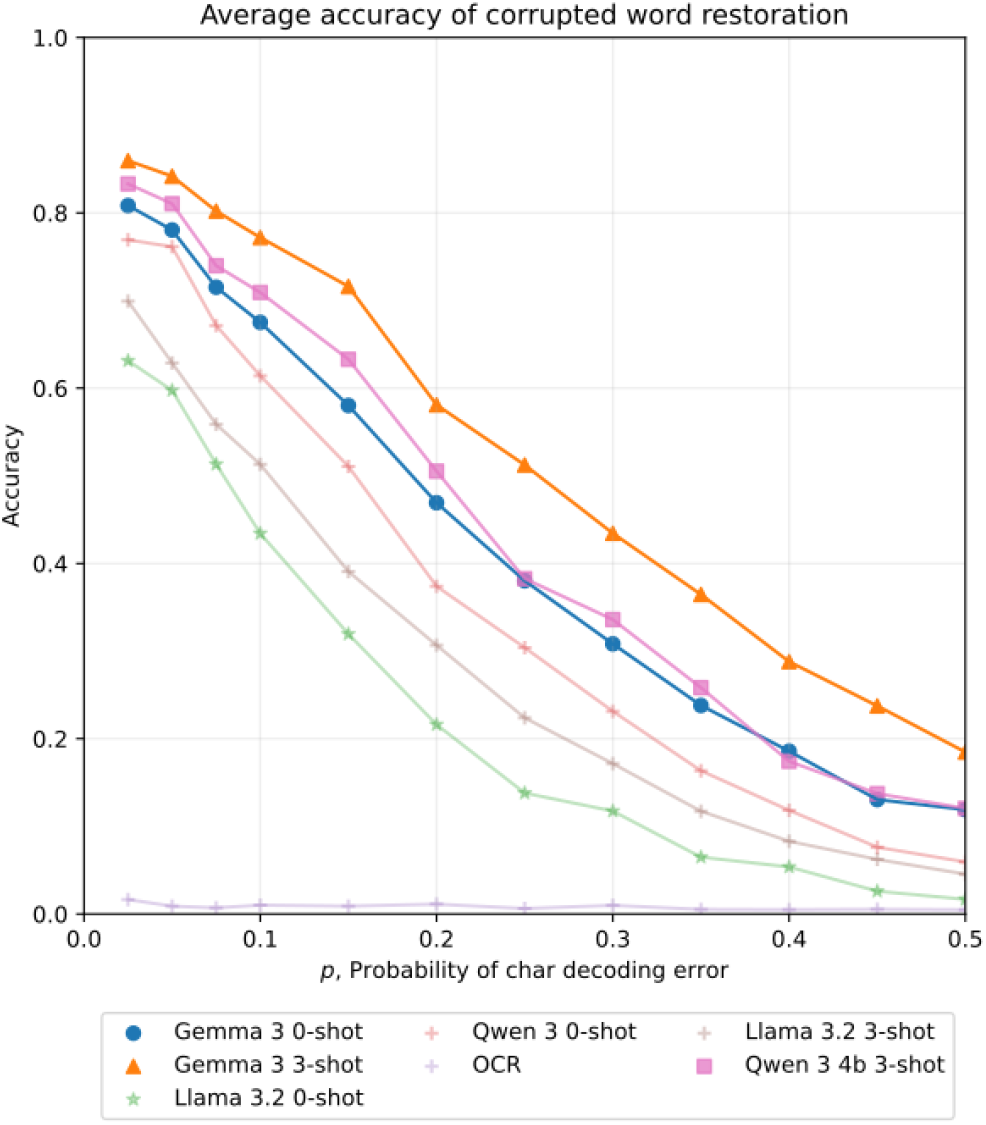
Average word restoration accuracy as a function spelling error probability for random errors. Curves for several LLMs are shown. (See key at the bottom.)

**Fig. 5.**
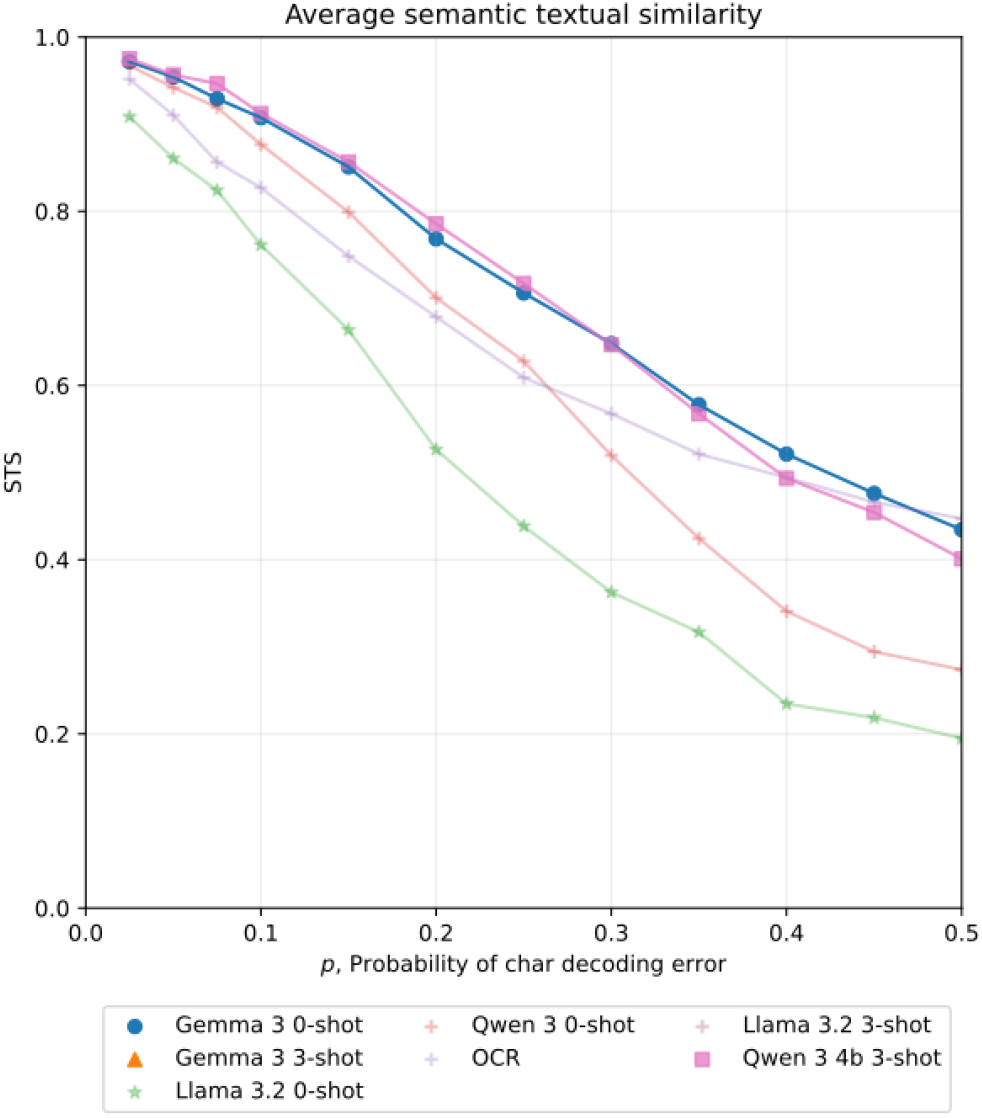
Average sematic textual similarity score (STS) for the reconstructed text as the function of spelling error probability for human-like errors. Curves for several LLMs are shown. (See key at the bottom.)

**Fig. 6.**
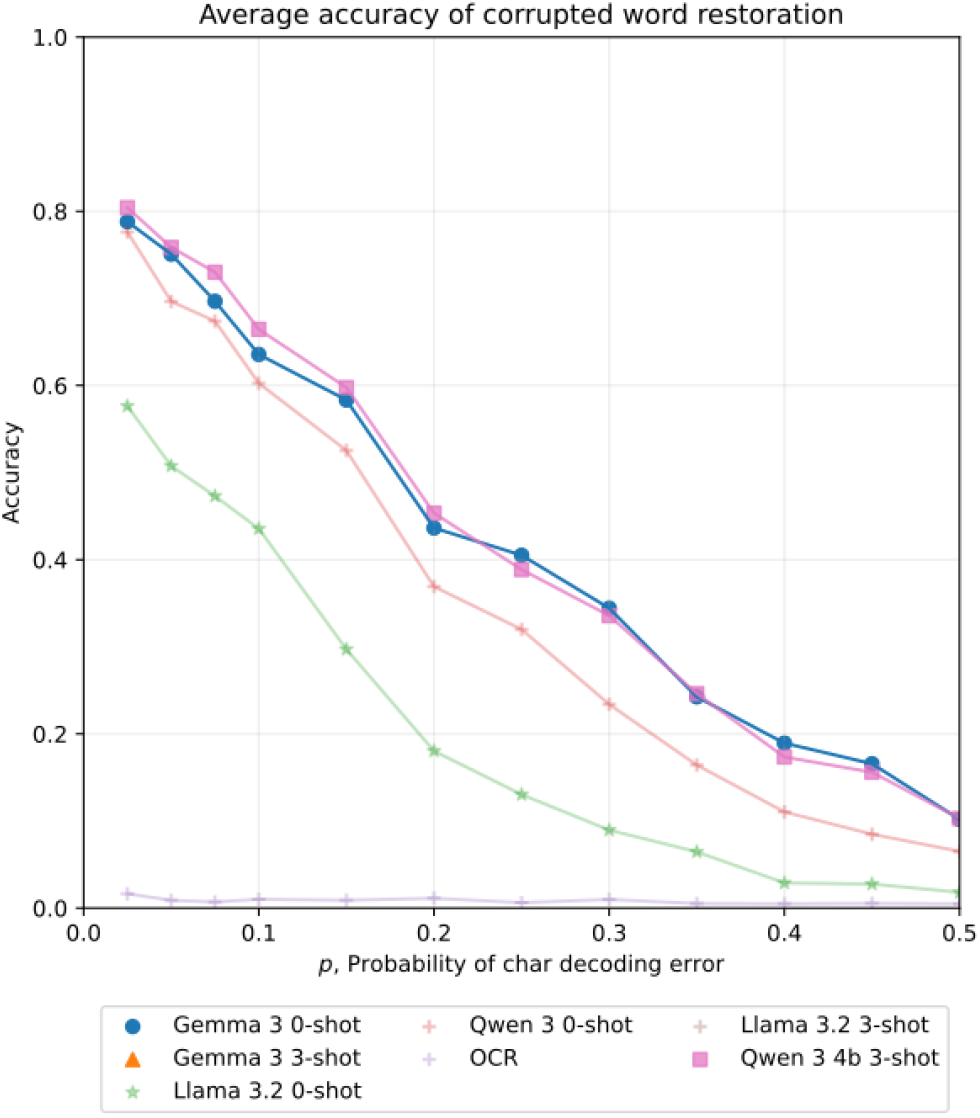
Average word restoration accuracy as a function spelling error probability for human-like errors. Curves for several LLMs are shown. (See key at the bottom.)

The Gemma 3 model demonstrated superior performance in both random and human-like settings compared to Qwen 3 and LLaMA 3.2. We attribute this difference to the tokenizer choice: Gemma 3 employs a SentencePiece tokenizer, whereas Qwen 3 and LLaMA 3.2 rely on Byte Pair Encoding (BPE). Prior studies suggested that SentencePiece tokenization is generally more effective for text correction tasks. In addition, the OCR fine-tuned model shows weaker performance, which can be explained by differences between the error distribution in OCR tasks and that of the corrupted text considered in this study.

Overall the difference in LLM performance was small between randomly generated errors and human-like errors. Since human-like errors are less homogeneous, they introduce additional complexity for the model. Nevertheless, in both settings LLM performance was strong.

## Discussion

Our study demonstrated that LLMs can be effective as post-processing components for P300 spellers, reconstructing text from corrupted outputs. This addresses one of the most persistent challenges in P300-based BCIs: the trade-off between communication speed and accuracy. By leveraging the contextual reasoning capabilities of modern LLMs, we showed that severely degraded speller output can be restored to coherent text, potentially enabling systems to operate with significantly fewer stimulus repetitions.

The consistent superiority of Gemma 3 across all experimental conditions warrants particular attention. With SentencePiece tokenization, Gemma 3 achieved the highest word restoration accuracy and semantic textual similarity scores under both random and human-like corruption patterns. This finding aligns with prior work suggesting that SentencePiece-based tokenizers are better suited for character-level text correction tasks (Kudo and Richardson, 2018), as they maintain greater flexibility in handling subword units and out-of-vocabulary characters. In contrast, models with BPE tokenizers were more prone to hallucinations as error probability increased, while Gemma 3 was not sensitive to this effect. Future work should focus on further investigation, including fine-tuning existing Gemma 3 models and adapting OCR-specific techniques to this task. Reducing inference time is also necessary.

The performance differential between error types was also instructive. Human-like errors, derived from empirical confusion matrices in row-column P300 paradigms (Gavett et al., 2012), were slightly more challenging than random substitutions. This is expected, as human-like errors exhibit systematic patterns that could conflict with the statistical distributions learned during pretraining.

The relatively poor performance of the OCR-fine-tuned ByT5 model, despite being designed specifically for character-level correction, highlights the importance of domain alignment in transfer learning. The error distributions in OCR tasks (Do et al., 2025), predominantly visual confusions such as “rn” for “m” or “cl” for “d”, from the substitution patterns observed in P300 spellers, which are driven by neural classification ambiguity rather than visual similarity.

The observed performance characteristics suggest a transformative pathway for P300 speller design. If an LLM can reliably correct text with up to 50% character-level errors, as our results indicate, system designers could substantially reduce the number of stimulus repetitions required for reliable signal acquisition. Users of P300 spellers often report cognitive fatigue from sustained attention to stimulus sequences. By reducing the required repetition count, LLM post-processing could alleviate this burden, making the system more sustainable for extended use. Additionally, the robustness of the approach could reduce the frustration associated with error correction, which requires users to navigate backward and reselect characters. Moreover, the near-real-time inference times (1.2–2.4 s per sentence) achieved on consumer-grade hardware suggest that LLM-based post-processing could be integrated without introducing unacceptable latency.

After an LLM-integrated P300 speller demonstrates reliable gains in real-time performance, the natural next step would be to leverage this enhanced capability to refine the feedback loop presented to the user during operation. As spelling trials accumulate and the system continuously processes incoming EEG data, dynamic feedback could be provided by visually highlighting keys or characters with the highest predicted probabilities of being the intended selection. Such visual cues would not only increase the transparency of the system’s internal state but also reduce uncertainty for the user. By embedding the LLM more deeply into this feedback mechanism, the system could improve these probability displays and offer context-aware suggestions such as next-character predictions and sentence completions. This richer, more intelligent feedback would boost user confidence in their selections, decrease the frequency of letter corrections, and ultimately improve both spelling speed and overall accuracy. As the LLM becomes increasingly attuned to the user’s intent and linguistic patterns, its function can evolve from that of error corrector to a more proactive conversational partner, one capable of anticipating user needs, handling ambiguous inputs, and supporting interactive dialogue. This progression would marks a fundamental shift from passive, unidirectional signal processing toward a collaborative human-AI interaction paradigm, where the system actively participates in the co-construction of meaning.

